# Tree bark scrape fungus: A potential source of laccase for application in bioremediation of non-textile dyes

**DOI:** 10.1101/2020.02.20.957365

**Authors:** H.M. Bhamare, R. Z. Sayyed, Najat Marraiki, Abdallah M. Elgorban, Asad Syed, Hesham Ali El-Enshasy

## Abstract

Although laccase has been recognized as a wonder molecule, and green enzyme, the use of low yielding fungal strains, poor production, purification, and low enzyme kinetics have hampered its larger-scale applications. Hence the present research was aimed to select high yielding fungal strains and to optimize the production, purification, and kinetics of laccase of *Aspergillus* sp. HB_RZ4. *Aspergillus* sp. HB_RZ4 produced a copious amount of laccase on under meso-acidophillic shaking conditions in a medium containing glucose and yeast extract. A 25 µM of CuSO_4_ enhanced the enzyme yield. The enzyme was best purified on Sephadex G-100 column. Purified enzyme resembled with the laccase of *A. flavus*. Kinetics of purified enzyme revealed the high substrate specificity and good velocity of reaction with ABTS as substrate. The enzyme was stable over a wide range of pH and temperature. The peptide structure of the purified enzyme resembled with the laccase of *A. kawachii* IFO 4308. The fungus decolorized various dyes independent of the requirement of a laccase mediator system (LMS). *Aspergillus* sp. HB_RZ4 came out as a potent natural producer of laccase, it decolorized the dyes even in absence of LMS and thus can be used for bioremediation.

## Introduction

Laccase (benzenediol: oxygen oxidoreductase, EC 1.10.3.2) belonging to a group of an enzyme called a multicopper blue oxidase (MCBO) exhibits wide substrate specificity ranging from phenolic to non-phenolic compounds [1]. It finds applications in various sectors such as biomedical [2], dye degradation [3] (paper industries for delignification [4-5], bioremediation [6], in biosensors [7] in cosmetic industry as melanin degraders [8] and as a enzymatic biofuel [9]. It is the major ligninolytic enzyme used in juice clarification [10]. Broad substrate promiscuity of laccase is the key perspective of laccase enzymology for various oxidoreductase reactions study. Laccase is a key biological mediator and best suitable alternative to chemical mediators hence regarded as a green enzyme in dye degradation which is the new era of dye degradation [11]. Synthetic dyes are broadly used in a wide range of industries including textiles, paper, printing, cosmetics, and pharmaceuticals. During dyeing, a 10–15 % of the dyes are lost in the effluent. Owing to their structural complexity most of these dyes resist biodecolorization. Bright colour, water-soluble, and acid dyes are highly resistant to the biodecolrization [12]. Although there are physic-chemical approaches available for removal of these dyes. However, these approaches are costly and not eco-friendly too [12].

High catalytic efficiency is another key feature of the enzyme in the bioremediation of dye effluent, sulphonamide, and other pollutants. This bioremediation is mediated by a laccase mediator system (LMS) [13]. Laccase has emerged as significant enzyme in mycoremediation of grey-water treatment as it substantially reduces the COD, BOD, and solids present in grey-water [14]. The new trend of forward osmosis with the help of laccase is used in micro-pollutant removal from wastewater and increasing potability of water [15]. Laccase is also majorly used in nitrogen organics biodegradation which is the major critical factor in water pollution with excellent catalytic performance and reusability [16]. It is the multifunctional lignin modifying enzyme (LME) having a great potential of bioremediation of dyes, organic pollutants and polycyclic compounds [17].

The laccase has a self as well as a cross-coupling mechanism in catalyzing single-electron oxidation playing an important role in removing non-degradable organic pollutants [18]. It is now used as effective and best alternative to chemical bleaching agents used for paper bleaching in the paper industry [19]. In order to further enhance the commercial potential and economically feasible uses of laccase, search for high yielding strains, optimization of physicochemical parameters for efficient production and factors regulating the laccase production need to be studied. Hence the present work was undertaken to search the best laccase producing fungus.

Nonetheless, high production cost and low efficiency of laccase has restricted its wider applications and compels the need to develop an economically feasible process [20]. A production yield of the enzyme depends on the type of producing strain as most natural strains are poor laccase producers. However, screening and selection of potent laccase producing fungi and the optimization of production conditions still appear as a crucial and vital approach of achieving high and cost-effective yields of laccase. There are reports on the improvement of laccase production through optimization of medium composition, and cultivation parameters [21].

## Materials and Methods

### Chemicals

All the chemicals used in present stud were purchased from Hi-media laboratories, Mumbai, India, Remazol Brilliant Blue R and 2,2’-azino-bis (3-ethylbenzothiazoline 6-sulphonic acid) (ABTS) were procured from Sigma Aldrich, USA.

### Source of culture

*Aspergillus* sp. HB_RZ4 obtained from the Department of Biotechnology, SSVPS’s Science College, Dhule, Maharashtra, India was used in present. It was previously isolated from tree bark scraping [22].

### Screening for laccase production

Three different media namely tannic acid agar (TAA) [23], guaiacol agar (GuA) and gallic acid agar (GAA) containing 0.5% tannic acid, 3% malt extract and 0.5% mycological peptone respectively were used to screen the production of ligninolytic enzymes. In GuA and GAA, tannic acid was replaced by guaiacol (0.01%) and gallic acid (0.5%) respectively. One plug (1 cm diameter) of *Aspergillus* sp. HB_RZ4 culture was grown on each plate at 32°C for 6 days and observed for the formation of brown halos around the fungal growth.

Alternatively one plug (one cm diameter) of *Aspergillus* sp. HB_RZ4 grown on selective basal media plates containing (gL^-1^): peptone, 3.0; glucose, 10.0; KH_2_PO_4_, 0.6; ZnSO_4_, 0.001; K_2_HPO_4_, 0.4; FeSO_4_, 0.0005; MnSO_4_, 0.05; MgSO_4_, 0.5; agar 2% supplemented with 0.1% (w/v) ABTS [24] at 32°C for 5 days.

### Production of laccase

For laccase production 2 plugs growth of fungus was grown in minimal medium (MM) containing (gL^-1^) glucose, 3.0; KH_2_PO_4_, 1.0, (NH_4_)2SO_4_, 0.26; MgSO_4_.7H_2_O, 0.5; CuSO_4_.7H_2_O, 00.5; 2,2-dimethyl succinic acid, 2.2; CaCl2.2H_2_O, 0.74; ZnSO4.7H_2_O, 0.6; FeSO_4_.7H_2_O, 0.5; MnSO_4_.4H_2_O, 0.5; CoCl_2_.6H_2_O, 0.1; vitamin solution 0.50 µl pH set to 4.5 [25] at 32°C for 12 days. The cell-free extract obtained following the centrifugation at 10,000 rpm for 15 minutes at 4°C was used for laccase assay with ABTS as standard (100-1000 µgmL^-1^) [26].

### Laccase assay

The reaction mixture comprising of 2.0 ml 100 mM sodium acetate buffer (pH 4.0), 80 µl ABTS and 20 µl of enzyme was incubated for 10 minutes [27] and oxidation of ABTS was recorded at 420 nm (εmax¬= 36000 M^-1^cm^-1^) and expressed as units per ml (UmL^-1^). One unit of the enzyme was defined as the enzyme required for the formation of 1 µM of product per min [28]. Micromole cation radical liberated per min per ml was calculated as follows [29].

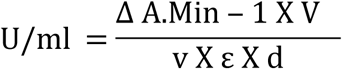

Where

V = Total reaction volume (mL)

v = Enzyme volume (mL)

ε = Extinction coefficient of ABTS at 420 nm (εmM) = 36 mM^-1^cm^-1^.

d = Light path of the cuvette (cm)

Δ*A*.min^-1^ = Absorbance change per minute at 420 nm.

### Estimation of fungal growth

Following the incubation, MM was filtered through Whatman filter paper No l, the resultant biomass was dried at 70°C and weighed till the constant weight and was expressed as mgmL^-1^.

### Optimization studies

Cultural conditions and media variables for optimum growth and laccase production were optimized by one Variable at a Time (OVAT) approach.

#### Influence of incubation period on growth and laccase production

In order to ascertain the exact time for optimum growth and laccase production, 2 plugs growth of 1cm diameter of fungus was grown in basal medium for 12 days at 32°C at 120 rpm (Kumar et al. 2016). Samples were withdrawn after every 24 h and were subjected for estimation of laccase activity and fungal growth.

#### Optimization of variables for growth and laccase production

The physical parameters viz. pH (2-10), temperature (20-55o C), time-course study (1-12 days) (Kumar et al. 2016)and nutrients such as carbon sources (@1.5%) such as glucose, sucrose, starch, maltose, lactose, fructose and glycerol, organic nitrogen sources (@1.5%) like L-asparagine, glutamic acid, glycine, L-proline, yeast extract, peptone, urea, inorganic nitrogen sources (@1.5%) such as NH_4_NO_3_, NaNO_3_, KNO_3_, NH_4_Cl, NH_4_H_2_PO_4_, and (NH_4_)2SO_4_, inducers (@10-50 µM) namely CuSO_4_, tween 80, veratryl alcohol, guaiacol, 2,5 xylidine, vanillic acid, gallic acid, ammonium tartrate and vanillin were optimized for optimum production of laccase [30].

### Purification of enzyme

The cell-free extract of production medium was precipitated with ammonium sulfate in the concentration range of 10 to 85% (w/v) under continuous stirring at 4°C. The precipitate was separated by centrifugation at 5,000×g for 10 min at 4°C and re-dissolved in 30 mL of sodium acetate buffer (100 mM, pH 4.5) and dialyzed with the same buffer by using Membrane filter No 110 with cut off 12-14 kDa (Hi-Media, Mumbai, India). The dialyzed fraction was loaded on DEAE-cellulose resin and eluted with a linear salt gradient (0-0.8 M NaCl) in sodium acetate buffer (100 mM, pH 4.5followed by further purification on Sephadex G-100 column. The active fractions were pooled and assayed for protein content and enzyme activity [12].

### Characterization of the purified enzyme

#### Determination of molecular weight of enzyme

The homogeneity and molecular weight of purified protein fraction were determined by Sodium Dodecyl Sulfate Polyacrylamide Gel Electrophoresis (SDS-PAGE). Purified fractions and standard protein marker (Ge-Nei, Bangaluru, India) were electrophoresed on SDS-PAGE comprising of resolving gel (10%) and stacking gel (5%) [31]. Followed the electrophoresis separated bands were stained with Coomassie Brilliant Blue R-250. The molecular mass was estimated by comparing the separated bands with standard protein markers. The protein content of supernatant at each stage was estimated as per Lowry et al. [32] bovine serum albumin (1000 µgmL^-1^) as a standard.

#### Determination of protein sequence of the enzyme by MALDI-TOF

The purified enzyme band obtained in SDS-PAGE gel was excised carefully and subjected to trypsin digestion [33]. The digested peptides were analyzed on the MALDI-TOF/TOF (Bruker Daltonics, Germany). Peptide mass fingerprint (PMF) analysis performed with Flex analysis software. The mass obtained in the PMF was submitted for MASCOT search in the database for identification of the protein and was compared with NCBI-nr database.

### Optimization studies on purified laccase

#### Influence of pH on enzyme activity and pH stability

The influence of pH on enzyme activity was investigated by dissolving the substrate (ABTS and guaiacol) in 50 mM of glycine-HCl buffer (pH 3.5), citrate phosphate buffer (pH 7.5) and glycine-NaOH buffer (pH 710). Following the incubation, at 34°C the enzyme activity was measured at 420 nm.

For studying the pH stability of enzyme the partially purified enzyme was pre-incubated at various pH ranges (2-10) for 60 min, 120 min. and 240 min. at 34°C followed by estimation of residual enzyme activity with ABTS substrate.

#### Influence of temperature on enzyme activity and thermal stability

The temperature profile of laccase activity was studied in 1.0 mM ABTS system. The oxidation of ABTS was carried out at a various temperatures in the range from 20-80°C [34]. For studying the thermal stability of the enzyme, the enzyme was incubated with 1.0 mM ABTS in the temperature range of 34-75°C for 150 min. Samples were withdrawn after every 30 min and subjected for estimation of enzyme activity.

#### Influence of inhibitors on laccase activity

Various inhibitors such as sodium azide (NaN3) (0.05-0.30 µmmL^-1^), cysteine (100-400 µm mL^-1^), EDTA (10-100 µmmL^-1^), halides (I-, Cl-, F-) (100-500 µm mL^-1^), thioglycolic acid (500-1500 µmmL^-1^) and thiourea (500-1500 µm mL^-1^) were evaluated to check their effect; partially purified enzyme was separately incubated with ABTS as a substrate and in different concentration of each inhibitor for 10 minutes at 34°C. The enzyme activity was determined.

#### Effect of activators (metal ions) on laccase activity

The effect of various metal ions such as Al^3+^, As^2+^, Ag^2+^, Cd^2+^, CO^2+^, Cu^2+^, FeCl_3_, FeSO_4_, Hg^2+^, Mg^2+^, Mn^2+^, MO^2+^, Ni^2+^, Zn^2+^, and Li^2+^ (1 mM) on laccase activity was examined by incubating the enzyme along with metal ions and 1.0 mM ABTS for 15 min at 34°C [35] followed by estimation of the residual activity of enzyme keeping reference enzyme as 100%.

#### Immobilization of enzyme

The immobilization of laccase was carried out by the entrapment method by using a 1:1 mixture of 1.5% (w/v) of gelatine and 3.0% (w/v) of sodium alginate. A 1.0 mL purified laccase was added to this mixture, mixed thoroughly for 10 min. at 25°C and withdrawn by using 5 mL sterile syringe with 22 gauge needle, the mixture was extruded into 2.0% (w/v) of pre-chilled CaCl_2_ solution to form beads of 2.0 to 3.0 mm diameter [36]. The immobilization efficiency was calculated by the following formula [37].

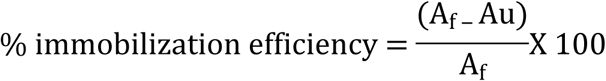

Where

A_f_= Specific activity of free enzyme

A_u_= specific activity of the unbound enzyme

### Enzyme kinetics

Kinetics studies of laccase were studied in ABTS (10-200 mM) as a substrate. The apparent Km and Vmax values were calculated by Michaelis-Menten and Lineweaver-Burk plot by using Graph Pad Prism 7.0 and Sigma Plot 12.0 software (San Diego, US).

### Evaluation of dye decolorization potential of enzyme

The ability of immobilized laccase to decolorize various non-textile dyes viz., methyl red, crystal violet, bromothymol blue, bromophenol blue, bromocresol purple, methylene blue, safranin, and methyl orange was examined in the presence of 2 mM of LMS (1-hydroxy benzotriazole [HBT]). The decolorization reaction mixture containing 50 mL 100 mM sodium acetate buffer (pH 4.5), dye (200 mgL^-1^) and enzyme (0.5 gm immobilized beads equivalent to 5 UmL^-1^) was incubated at 34°C for 96 h [12]. The degree of dye decolorization was estimated by recording the change in absorbance at a respective λmax and calculated as percent decolorization by using the following formula [37].

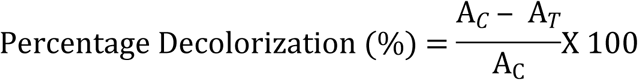

Where,

A_C_ is the absorbance of control

A_T_ is the absorbance of the test sample.

### Statistical analysis

All experiments were carried out in triplicate and the results were expressed as mean ± standard error/deviation.

## Result and Discussion

### Screening and production of laccase

*Aspergillus* sp. HB_RZ4 produced brown halos around and under the growth on GuA plate indicating the production of ligninolytic enzymes. It oxidized ABTS from ABTS agar and produced a green halo around the mycelia growth confirming the production of laccase. During submerged fermentation under shaking conditions at 32°C on 8th day of incubation *Aspergillus* sp. HB_RZ4 produced 6.22 UmL^-1^ laccase.

### Optimization of laccase production

#### Influence of incubation period on laccase activity

In the time-course studies, maximum laccase activity (8.422 UmL^-1^) and optimum growth (0.0021 mgmL^-1^) were evident on the 8th day of incubation (Figure 1).

**Figure 1.**
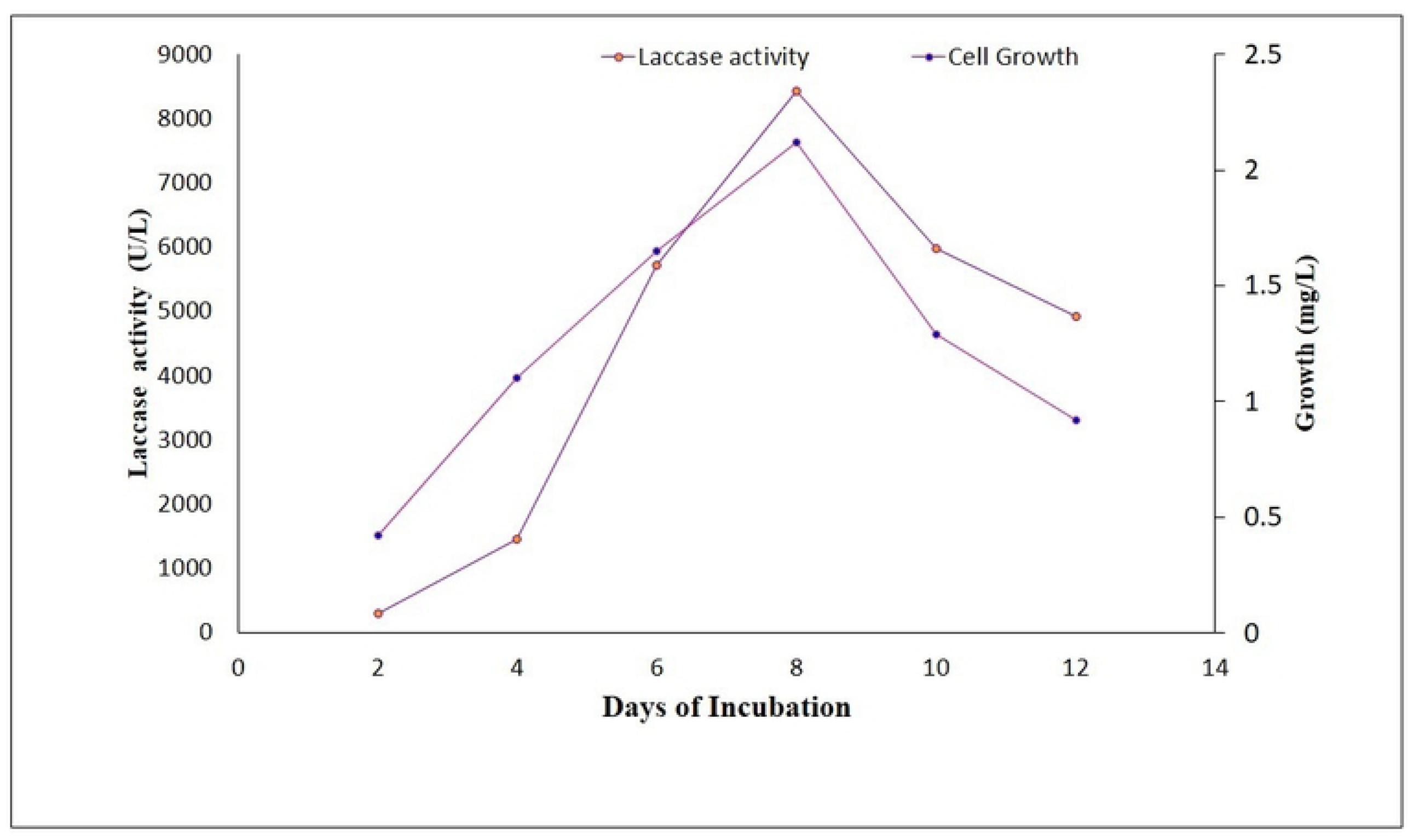
Influence of incubation period on laccase production in basal medium. Samples were withdrawn after every 24 h and were subjected for estimation of laccase activity and fungal growth.

#### Influence of pH and temperature on laccase production

The pH and temperature profile of laccase activity revealed that optimum enzyme activity (8.125 UmL^-1^) was obtained at an acidic pH value of 4.5 and incubation temperature of 34°C. pH values below and above 4.5 affected the enzyme activity, similarly incubation temperature below or above 34°C also affected the growth as well as laccase production.

#### Influence of carbon and nitrogen source on laccase production

Among the various carbon sources used for the production of laccase in *Aspergillus* sp. HB_RZ4, glucose resulted in a 12.45 fold increase in laccase production. Glycerol containing media resulted in the lowest laccase yield (2.761 UmL^-1^). The order of carbon sources supporting laccase production was glucose> sucrose> starch> maltose> lactose> fructose> glycerol. Among the organic and inorganic nitrogen sources, maximum laccase activity (6.581 UmL^-1^, 11.7 fold increase) was obtained with yeast extract while NH4NO3 served as the best inorganic nitrogen source as it yielded laccase activity of 3.97 UmL^-1^ (Table 1).

**Table 1.** Influence of nitrogen sources on the production of laccase in *Aspergillus* sp. HB_RZ4.

| Organic<br>nitrogen<br>source | Laccase production (UL<sup>-1</sup>) |  | Specific<br>activity<br>(Umg<sup>-1</sup>) | Fold<br>increase |
| --- | --- | --- | --- | --- |
|  | In absence<br>of inducer | In presence<br>of inducer |  |  |
| L-asparagine | 2.55 | 1.225 | 30.4 | 4.79 |
| Glutamic acid | 3.96 | 1.949 | 32.7 | 4.92 |
| Glycine | 3.84 | 1.087 | 26.3 | 2.83 |
| L-proline | 1.58 | 3.968 | 78.3 | 2.51 |
| Yeast extract | 5.62 | 6.581 | 208.8 | 11.7 |
| Peptone | 4.12 | 3.951 | 78.9 | 9.37 |
| Urea | 1.36 | 4.174 | 18.5 | 2.32 |
| NH<sub>4</sub>NO<sub>3</sub> | 3.97 | 2.649 | 64.0 | 6.67 |
| NaNO<sub>3</sub> | 3.02 | 1.492 | 32.5 | 4.93 |
| KNO<sub>3</sub> | 1.69 | 2.080 | 11.2 | 1.23 |
| NH<sub>4</sub>Cl | 2.08 | 0.996 | 25.7 | 4.78 |
| NH<sub>4</sub>H<sub>2</sub>PO<sub>4</sub> | 3.09 | 1.277 | 31.6 | 4.13 |
| (NH<sub>4</sub>)<sub>2</sub>SO<sub>4</sub> | 2.15 | 0.9621 | 23.2 | 4.47 |

| Organic nitrogen source | Laccase production (UL<sup>-1</sup>) |  | Specific activity (Umg<sup>-1</sup>) | Fold increase |
| --- | --- | --- | --- | --- |
|  | In absence of inducer | In presence of inducer |  |  |
| L-asparagine | 2.55 | 1.225 | 30.4 | 4.79 |
| Glutamic acid | 3.96 | 1.949 | 32.7 | 4.92 |
| Glycine | 3.84 | 1.087 | 26.3 | 2.83 |
| L-proline | 1.58 | 3.968 | 78.3 | 2.51 |
| Yeast extract | 5.62 | 6.581 | 208.8 | 11.7 |
| Peptone | 4.12 | 3.951 | 78.9 | 9.37 |
| Urea | 1.36 | 4.174 | 18.5 | 2.32 |
| NH<sub>4</sub>NO<sub>3</sub> | 3.97 | 2.649 | 64.0 | 6.67 |
| NaNO<sub>3</sub> | 3.02 | 1.492 | 32.5 | 4.93 |
| KNO<sub>3</sub> | 1.69 | 2.080 | 11.2 | 1.23 |
| NH<sub>4</sub>Cl | 2.08 | 0.996 | 25.7 | 4.78 |
| NH<sub>4</sub>H<sub>2</sub>PO<sub>4</sub> | 3.09 | 1.277 | 31.6 | 4.13 |
| (NH<sub>4</sub>)<sub>2</sub>SO<sub>4</sub> | 2.15 | 0.9621 | 23.2 | 4.47 |

#### Purification of laccase

Among the various methods used for purification, the best laccase of *Aspergillus* sp. HB_RZ4 was obtained on the Sephadex G-100 column. This method yielded total protein content of 2.0 mg, enzyme activity of 1.212 U and specific activity of 465 Umg^-1^ proteins giving rise to 6.21% purification yield with 65 fold purification. Salt precipitation and DEAE-cellulose method resulted in minimum protein content (0.2 and 0.7 mg); low enzyme activity (93 and 105 U), less specific activity (60 and 150 Umg^-1^) and poor purification yield (4.73 %) and minimum (8.5 and 21) fold purification (Table 2).

### Characterization of enzyme

#### Determination of molecular weight

The molecular mass of the purified laccase fraction as obtained from SDS-PAGE was approximately 62 kD (Figure 2).

**Figure 2.**
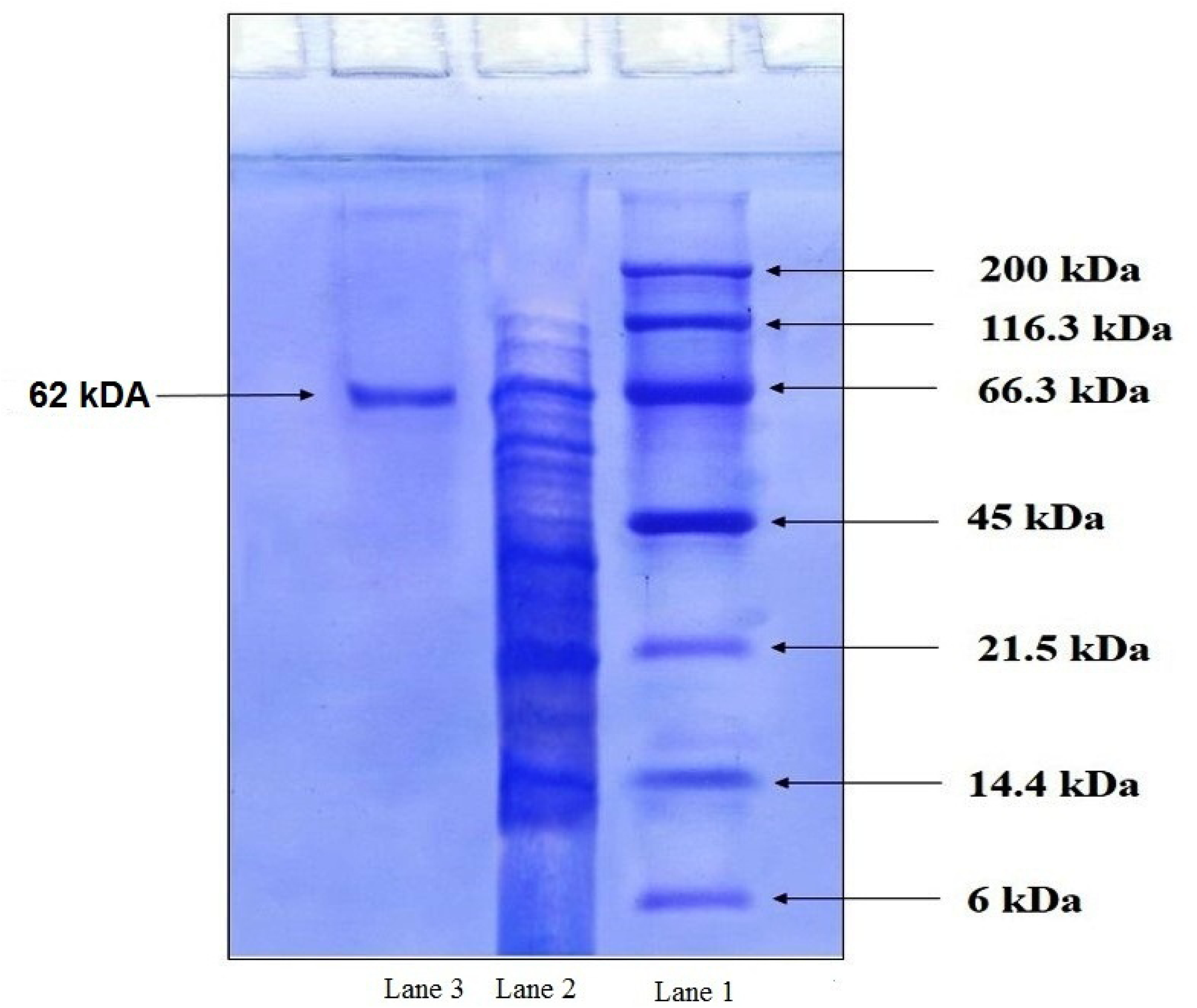
SDS-PAGE analysis for the molecular mass of protein of *Aspergillus* sp. HB_RZ4 Purified fractions of laccase (Lane 2) and standard protein marker (Lane 1) were electrophoressed on SDS-PAGE followed by staining with Coomassie Brillient Blue R-250 and the molecular mass of purified proteins was estimated by comparing it with the standard protein markers.

#### Determination of protein sequence of the enzyme by MALDI-TOF

Among the various trypsin digested peptide fragments; 10 peptides were hit in the protein database by Mascot peptide mass fingerprinting search engine. The amino acid sequences of each peptide of laccase exhibited a significant mascot score of 75 and p-value < 0.05 (Figure 3) with a known sequence of NCBI: GAA87354.1. These 10 peptides corresponded to 29.33% sequence coverage and showed homology with the laccase of *Aspergillus kawachii* IFO 4308 (NCBI: GAA87354.1).

**Figure 3.**
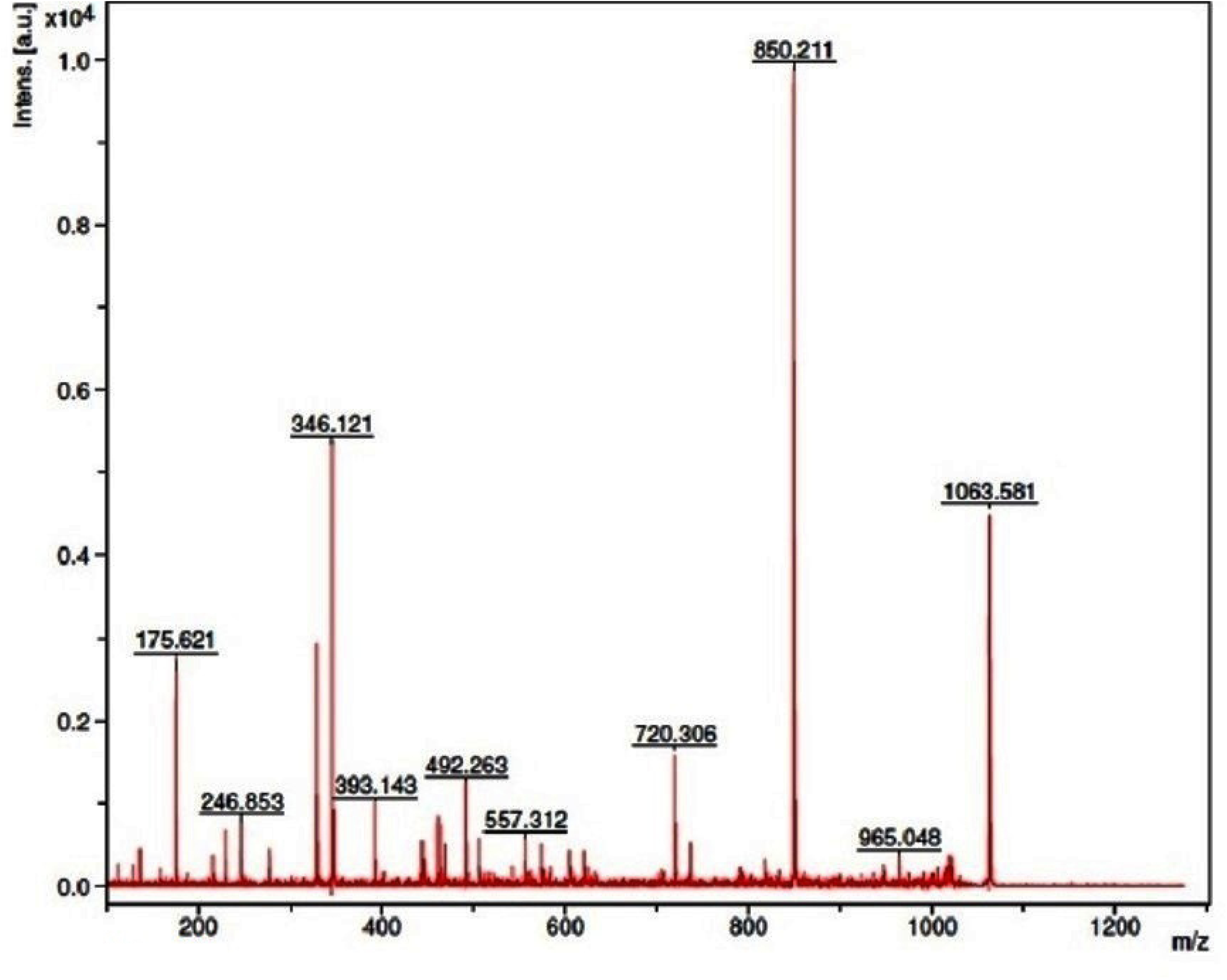
MALDI-TOF mass spectrum of trypsin digested peptide map of laccase. The purified enzyme band obtained in SDS-PAGE was digested by trypsin and subjected for peptide mass fingerprint analysis by using Flex analysis software and MASCOT search in the database and peptide/proteins were compared with NCBI-nr database.

### Optimization studies with purified enzyme

#### Influence of pH on laccase activity and stability

The purified laccase of *Aspergillus* sp HB_RZ4 showed pH optima of 4.5 with 100 % and pH optima of 6.0 with 99.9% residual activity for ABTS and guaiacol respectively. At these pH values the % residual activity was 100% and 99.9 % respectively. The enzyme was stable over the range of pH (neutral to alkaline) for a longer period i.e. 120 and 240 h; higher stability was evident at pH neutral pH.

#### Influence of temperature on laccase activity and stability

The enzyme was active over the wide range (20°C to 60°C) of temperature, 34°C being the optimum temperature till 90 min. of incubation. Increasing temperature beyond 34°C and incubation period of more than 90 min affected the enzyme activity. Good stability (97%) of the enzyme was obtained at 34°C after 90 minutes. Further increase in temperature above 34°C and prolonged incubation period beyond 90 min, affected the enzyme activity.

#### Influence of inhibitors and activator (metal ions) on laccase activity

Studies on the influence of different concentrations of inhibitors and metal ions revealed that some inhibitors affect the enzyme activity even at lower concentrations while others did not affect even in relatively higher concentrations. Sodium azide completely inhibited the enzyme activity at 0.30 µmmL^-1^ whereas L-cysteine even at higher concentration (400 µmmL^-1^) did not affect the enzyme activity. Halides strongly inhibited the enzyme; fluoride caused 98.72% inhibition even at low (25 µmmL^-1^) concentration. Chloride, bromide, and iodide at (300 µmmL^-1^) caused 96.12 %, 94%, and 94.12% inhibition respectively. Thioglycolic acid produced strong inhibition (97.10%) than thiourea (91.55%).

Other metal ions like Al3+, As2+, Cd2+, CO2+, and Li2+ significantly inhibited the activity of enzyme, Ag2+, Hg, FeSO4, and FeCl3 showed 90, 95, 78 and 76 % inhibition respectively. While Cu^2+^, Mo^2+^, Mn^2+^, Zn^2+^ enhanced the enzyme activity. The presence of Cu^2+^ significantly boosted the enzyme activity 8.125 UmL^-1^ to 8.692 UmL^-1^ (Figure 4) whereas vanillin, ammonium tartrate gallic acid and vanillic acid failed to enhance enzyme on the contrary they affected the activity of the enzyme. A 25 µM of CuSO_4_ was a threshold level for optimum laccase activity and fungal growth (8.692 UmL^-1^, 0.019 mgmL^-1^).

**Figure 4.**
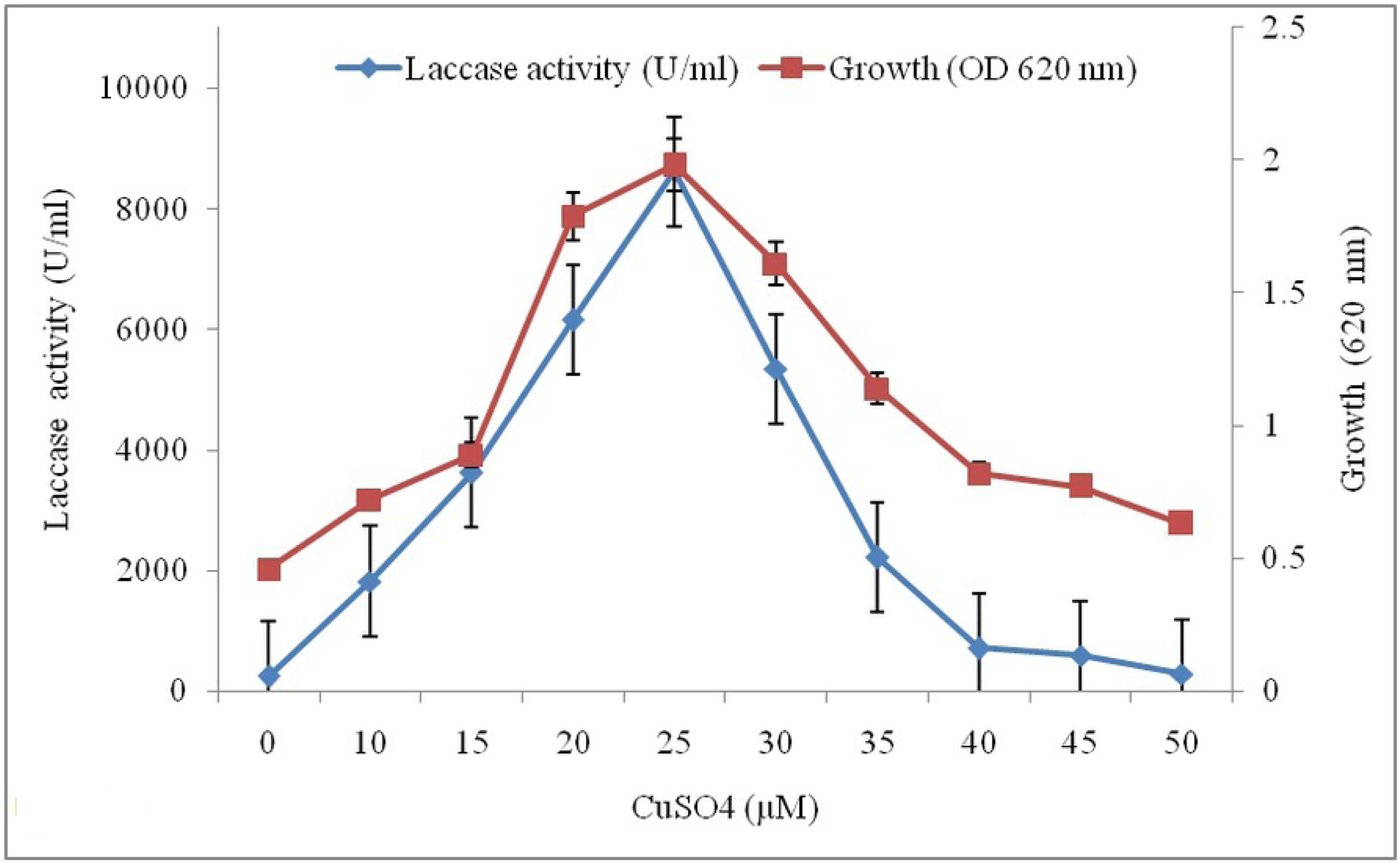
Influence of various concentration of CuSO_4_ on laccase activity. The reaction mixture contained the enzyme along with CuSO_4_ (0-50 µM) for 15 min at 34°C. The enzyme activity was measured with 1.0 mM ABTS by keeping reference enzyme as 100%.

#### Evaluation of dye decolorization potential of enzyme

LMS had negligible effects on the dye decolorization potential of laccase, absence of LMS 86%, 92, 98% and 95% decolorization of the bromothymol blue, methyl red, bromophenol blue, and bromocresol purple respectively was obtained. On the contrary, the presence of LMS inhibited the decolorization of methyl red, safranin, and methyl orange. But in the case of congo red, crystal violet and methylene blue LMS increased the decolorization of these dyes by 1.49, 1.99 and 3.47 fold respectively.

### Enzyme immobilization

The immobilized enzyme exhibited 92% enzyme efficiency (8.556 UmL^-1^) vis-à-vis 100% efficiency (9.30 UmL^-1^) of the free enzyme at and it deteriorated with the increasing period, on 8^th^ day of incubation enzyme efficiency fall down to 48.38% (4.5 U mL^-1^).

### Enzyme kinetics

The kinetic parameters Km and Vmax of purified laccase were 26.8 mM and 7132.6 mM min^-1^ respectively.

## Discussion

Fungi are the most widespread saprophytes that degrade the organic matter through the secretion of a number of lignolytic enzymes including laccase. Formation of brown halos around and under the growth of *Aspergillus* sp. HB_RZ4 on GuA plate is due to the oxidation of guaiacol indicating the production of lignolytic enzymes. The formation of green halo around the mycelia growth is due to the oxidation of ABTS (substrate) to a stable colored product, 2,2-azinobis-(3-ethylbenzothiazoline-6-sulfonate) under the influence of laccase [38]. Since ABTS is a specific substrate for laccase, its oxidation indicates that the enzyme produced by *Aspergillus* sp. HB_RZ4 is a true laccase [24]. Both these screening tests confirmed the ability of *Aspergillus* sp. HB_RZ4 to produce laccase. It produced 6.22 UmL^-1^ laccase on the 8th day of incubation during shake flask growth at 32°C. Ghosh and Ghosh [39] have reported more laccase production from *A. flavus* on the 20th day of incubation. Kumar et al. [24] have reported optimum laccase production (17.39 IUmL^-1^) production in *A. flavus* on the 12th day of incubation. Some fungal species require a longer production time i.e. 12–30 days [40]. Sivakumar et al. [41] reported a 4.60 IU mL^-1^ laccase yield after 12 days of incubation under static conditions. In many fungi, laccase synthesis is activated by type and nature of carbon or nitrogen source which determines the duration of the production cycle. Therefore, the best laccase producing organism should produce high yields of laccase in a short fermentation cycle [42]. A higher yield of laccase in less time (8 days) reflects the metabolic efficiency of organisms and suggests the possibility of exploitation of organism for cost-effective production of laccase at commercial scale. The optimization of physicochemical parameters boosted the enzyme yield. Optimum laccase yield in glucose and yeast extract containing medium is due to the rapid assimilation of glucose as it is readily oxidizable sugar and yeast extract is the source of all amino acids required for the synthesis of laccase [43]. Senthivelan et al. [44] have reported the production of laccase in *Penicillium chrysogenum*. The statistical optimization enhanced the enzyme activity to 7.9 U mL^-1^ against 6.0 U mL^-1^ obtained under un-optimized conditions.. Laccase production in many fungi including *P. chrysogenum* has been reported acidic pH and at mesophilic temperature. Media composition, presence or absence of metal ions and types and levels of nutrients are known to regulate the expression of laccase isozyme genes. The effects of organic compounds on laccase production depend on the compound structure, fungal strain, and growth stage [45].Fewer purification yields with ammonium sulfate precipitation may be due to the denaturation of the enzyme by ammonium sulfate. No purification with DEAE cellulose may be due to the ability of enzymes to absorption on the cellulose matrix. Good purification yields with the Sephadex G-100 column can be attributed to the better adsorption of enzyme on Sephadex gel. Many fungal laccases have been purified using Sephadex G-100 resins. Kumar et al. [24] have reported the purification of laccase of *A.flavus* on Sephadex G-100 resin. Patel and Gupte [46] have reported purification of laccase of *Trichoderma giganteum* AGHP using Sephadex G-75 and have reported an enzyme yield of 10.49 % with 3.33 fold purification. A 70-fold purification of laccase from *Stereum ostrea* using ammonium sulfate precipitation followed by Sephadex G-100 column chromatography has been reported by Vishwanath et al. [47]. The molecular weight of purified laccase of *Aspergillus* sp. HB_RZ4 was 62 k as evident in SDS-PAGE gel stained by coomassie brilliant blue (Fig. 3). The molecular weight of the fungus resembled with the molecular weight of laccases reported in other white-rot fungi [48]. Patel and Gupte [46] reported the molecular weight of 66 kDa by SDS-PAGE. The laccase was purified by Sephadex G-100 gives good specific activity o which is better than laccase from *Trametes versicolor* [49].Good activity and stability of enzyme for longer periods at wider pH range (acidic to alkaline) and a broad range of temperature (20°C to 60°C) is due to the presence of specific substrates like ABTS and guaiacol. Kumar et al. [24] have reported good laccase activity over the range of pH and temperature in *A. flavus*. The inhibitory effect of sodium azide, cysteine, EDTA, halogens, thioglycolic acid and thiourea on laccase activity was studied. The drastic decrease in enzyme activity is due to a change in pH of the medium that produces exert an inhibitory action on enzyme activity [50]. Severe reduction in enzyme activity by sodium azide and cysteine is due to the reason that sodium azide binds to the copper site of the enzyme and blocks the internal electron transfer reaction. Laccases are sensitive for metals even at low concentrations and inhibit the laccase activity. Good induction of enzyme activity in presence of CuSO4 is due to the filling of type-2 copper-binding sites with copper ions and due to the reason that Cu is the major inducer for laccase [51] as the catalytic center of the enzyme contains Cu ions. Xin and Geng [52] also copper sulfate best inducer for laccase production in *Trametes versicolor*. Mann et al. [53] reported 0.75 and 0.4 mM concentration of copper as the best level to induce laccase production in Ganoderma lucidum. Inhibition of laccase activity above 25 µM of CuSO_4_ may be because a higher concentration of copper inhibits the growth of fungi [54]. Potent inhibition of enzyme activity by Ag^2+^ and Hg^2+^ can be attributed to the formation of SH groups with enzyme leading to the inactivation of enzyme moreover these enzymes are known to possess antimicrobial activity [40]. This interaction of enzyme with metals has great significance for the better understanding and development of a process for bioremediation of xenobiotics, textile dyes, and grey-water. The effect of metal ions on laccase activity was based on the type of metals used since metal ions greatly influence the catalytic activity of the enzyme. The activation or inhibition of enzymes also regulates the turnover rate of enzymes. The enzyme was able to decolorize roughly 88%, 96% and 99% bromothymol blue, bromocresol purple and bromophenol blue respectively in presence of HBT. Copete et al [55] have reported 15% to 40% decolorization of various dyes by laccase producing *Leptosphaerulina* sp. and have found enhanced decolorization in the presence of a mediator. Zuo et al. [13] have reported 84.9% decolorization of bromocresol by *Pleurotus ostreatus* HAUCC 162 and have claimed the involvement of the effect of mediator HBT in increasing decolorization. Enzymes immobilization offers the reuse of enzyme, more stability, and resistance under diverse conditions and improves the catalytic activity of laccases [24]. Fungal laccases typically have 3 to 10 glycosylation sites, and 10 to 50% of their molecular weight is attributed to glycosylation and deglycosylation of laccase affects its enzyme kinetics [56].After SDS-PAGE electrophoresis a band excised from the gel was subjected for identification by using MALDI_TOF analysis. A band at ∼ 62 kDa was digested with trypsin into 10 fragments in the range of P1 to P10 (22 to 250 amino acid sequence) (Table 3a and 3b). Mascot database exhibited the 29.33% resemblance with laccase of *A. kawachii* IFO 4308 (NCBI: GAA87354.1) and confirmed that the purified enzyme was laccasev[57]. Km and Vmax of purified laccase were 26.8 mM which is a good activity of the enzyme. Tinoco et al. [58] have reported the Km values in the range 8 to 79 µmol for ABTS with different strains of *Pleurotus ostreatus.*

**Table 3a.**
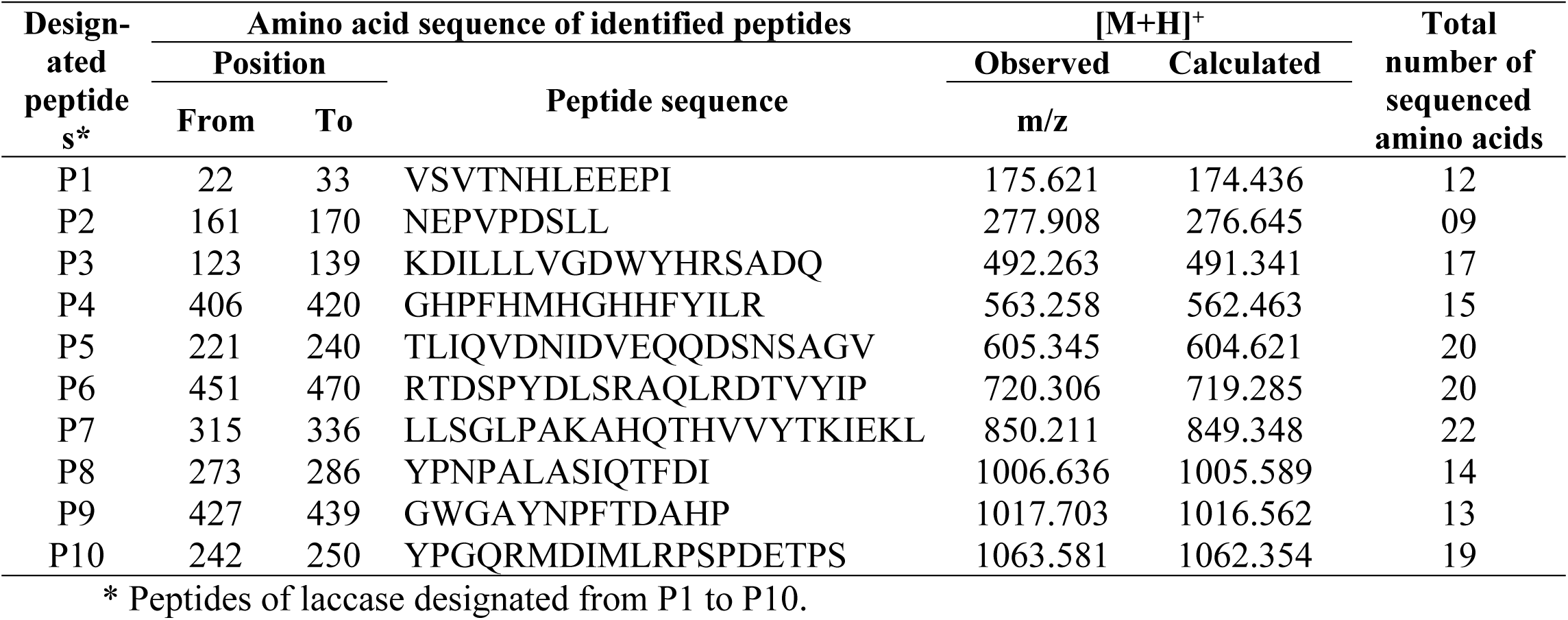
Peptide ions of trypsin digest of laccase of *Aspergillus* sp.HB_RZ4.

**Table 3b.** Amino acid content of sequenced peptides of laccase obtained by trypsin digestion.

| Amino acid | Number of amino acids per peptide of hyaluronidase* |  |  |  |  |  |  |  |  |  |  |
| --- | --- | --- | --- | --- | --- | --- | --- | --- | --- | --- | --- |
|  | P1 | P2 | P3 | P4 | P5 | P6 | P7 | P8 | P9 | P10 | Total |
| Gly (G) | - | - | 1 | 2 | 1 | - | 1 | - | 2 | 1 | 08 |
| Ala (A) | - | - | 1 | - | 1 | 1 | 2 | 2 | 2 | - | 09 |
| Val (V) | 2 | 1 | 1 | - | 3 | 1 | 2 | - | - | - | 10 |
| Leu (L) | 1 | 2 | 3 | 1 | 1 | 2 | 4 | 1 | - | 1 | 16 |
| Ile (I) | 1 | - | 1 | 1 | 2 | 1 | 1 | 2 | - | 1 | 10 |
| Ser (S) | 1 | 1 | 1 | - | 2 | 2 | 1 | 1 | - | 2 | 11 |
| Thr (T) | 1 | - | - | - | 1 | 2 | 2 | 1 | 1 | 1 | 09 |
| Cys (C) | - | - | - | - | - | - | - | - | - | - | 00 |
| Met (M) | - | - | - | 1 | - | - | - | - | - | 2 | 03 |
| Asp (D) | - | 1 | 3 | - | 3 | 3 | - | 1 | 1 | 2 | 14 |
| Asn (N) | 1 | 1 | - | - | 2 | - | - | 1 | 1 | - | 06 |
| Glu (E) | 3 | 1 | - | - | 1 | - | 1 | - | - | 1 | 07 |
| Gln (Q) | - | - | 1 | - | 3 | 1 | 1 | 1 | - | 1 | 08 |
| Phe (F) | - | - | w- | 2 | - | - | - | 1 | 1 | - | 04 |
| Tyr (Y) | - | - | 1 | 1 | - | 2 | 1 | 1 | 1 | 1 | 08 |
| Trp (W) | - | - | 1 | - | - | - | - | - | 1 | - | 02 |
| Lys (K) | - | - | 1 | - | - | - | 3 | - | - | - | 04 |
| Arg (R) | - | - | 1 | 1 | - | 3 | - | - | - | 2 | 07 |
| His (H) | 1 | - | 1 | 5 | - | - | 2 | - | 1 | - | 10 |
| Pro (P) | 1 | 2 | - | 1 | - | 2 | 1 | 2 | 2 | 4 | 15 |
| **Total** | **12** | **09** | **17** | **15** | **20** | **20** | **22** | **14** | **13** | **19** | **161** |

## Conclusion

In the present study, Aspergillus sp. HB_RZ4 produced copious amounts of the extracellular laccase in MM under mesophilic conditions at acidic pH. Its production was not affected by many of the conventional inhibitors and chemicals. The stability of laccase over the range of pH and temperature and ability to decolorize the dye without requiring LMS makes it a magic molecule for cost-effective production of it and its exploitation in bioremediation of effluent containing dyes.

## Acknowledgement

The Researchers Supporting Project Number (RSP-2019/56) King Saud University, Riyadh, Saudi Arabia and MOE and UTM-RMC, HICOE, Malaysia for funding this research through the Research Group Project No. R.J130000.7846.4J262.

